# Plasmid carriage and the unorthodox use of fitness in microbiology

**DOI:** 10.1101/810259

**Authors:** Carlos Reding

**Affiliations:** Experimental Medicine Division, Nuffield Department of Medicine, University of Oxford.UK

## Abstract

The link between fitness and reproduction rate is a central tenet in microbiology, and indeed evolutionary biology: Mutants reproducing faster than the dominant wild-type are favoured by selection, but otherwise the mutation is lost. This link was given by Ronald Fisher in 1930 under the assumption that fitness can only change through mutations that boost or hinder growth rate, whence the use of logarithms on growth data by experimentalists. Here I show that logarithms are highly sensitive to sampling times, resulting in fitness estimates that are not constant over the growth of bacterial cultures. This variability invalidates typical selection measurements, and ‘unfit’ mutants can be co-maintained if they reach their equilibrium. And this is what I observed in competition assays between two *Escherichia coli* constructs, one of which harbours a non-transmissible plasmid that protects against tetracycline (pGW155B), without using the antibiotic. Despite growing 40% slower than its drug-sensitive counterpart, the construct harbouring the plasmid persisted throughout the competition. And, perhaps more importantly, maintained the plasmid. My study suggests that reliance on growth rate masks that selection on plasmid carriage may be stronger than previously thought—explaining the seemingly-paradoxical abundance of plasmids in nature.

## I. INTRODUCTION

Population genetics [1–4] and microbiology [5, 6] textbooks define the fitness of microbial populations as their intrinsic growth rate. This notion stems from the use of the exponential growth model to study natural selection, where fitness (*m*) is given by the exponential model *N*(*t*) = *N*_0_*e*^*mt*^—originally introduced by Fisher [7] as the Malthusian parameter, whence *m*. Here *N*_0_ is the initial number of individuals of a given genotype, and *N*(*t*) the number of individuals at time *t*. The ratio *W*_*ij*_ = *m*_*i*_/*m*_*j*_ is the gold-standard [8] metric to estimate the fitness of one genotype with respect to another (*W*_*ij*_). Strength of selection is then measured as *S*_*ij*_ = (*m*_*i*_ − *m*_*j*_)/*m*_*j*_ = *W*_*ij*_ − 1 [1, 8] where *S*_*ij*_ is the selection coefficient. This approach is an over-simplification [9] since microbial growth is density-dependent, and yet, it still is heavily used by evolutionary [8, 10–16] and clinical [17–20] microbiologists alike by using logarithms on growth data to estimate the Malthusian parameter *m* [8, 14, 15, 18, 19]. Some studies do use a logistic growth model [20], but relative fitness is then reduced to differences in the growth rate parameter alone. Despite its mathematical simplicity, the use of the exponential growth model is said [10] to aggregate the contribution of each growth phase to fitness. But this is not reflected in the exponential growth model with only one parameter. Thus, others [21, 22] tried to model the contribution of each of these phases to fitness. Alas, intrinsic growth rate remains as central to these models as it is for the exponential growth model.

If fitness—as intrinsic growth rate—is an inherited trait, it is reasonable to expect that mutants with greater fitness will have greater representation in the next generation eventually replacing the wild-type genotype [1, 7, 23]. This rationale underpins the study of plasmid maintenance in bacteria. Curiously, despite its central place in plasmid biology, this rationale cannot explain the abundance of plasmids in nature [24–27]. Plasmids are extra-chromosomal elements that bacteria can acquire through horizontal gene transfer to thrive in unfavourable environments [24]. In favourable environments however, where plasmids do not add an advantage, they impose a fitness cost derived from the synthesis of additional genetic material during the replication and expression of plasmid-borne genes [28, 29]. Thus, conventional wisdom holds that plasmids should not exist without positive selection [24]—indeed a typical assumption also used in epidemiological models [30, 31]. But they do, and are ubiquitous in nature [32], creating a so-called *plasmid paradox* [24, 26]. How can we explain this paradox? There is evidence showing that plasmids can undergo horizontal-gene transfer fast enough to persist despite the costs [33]. The problem here is that most of the known plasmids are *not* transmissible [32]. So, how can we explain their abundance? The question then is whether a myriad of conditions exist to select for plasmid-carriage, or the logic above behind the notion of fitness is sub-optimal. After all, the availability of data testing the predictions of population genetics theory is extremely scarce [34–36]—whether plasmids do indeed incur fitness costs at all has been questioned in the past [11, 37].

The fitness metric *W*_*ij*_ above implies that the ratio *m*_*i*_/*m*_*j*_ is a constant, following the view that fitness is inheritable and therefore it can only change through mutations that affect *m* [8]. This is mathematically consistent since the Malthusian parameter *m* is also a constant, but the ratio is rarely checked experimentally. I did, and below I show that, rather, *m*_*i*_/*m*_*j*_ changes predictably through time—not just in evolutionary timescales. This means that relative fitness depends not only on *m*, but *also* on when it is measured. Following this logic, in this study, I found conditions to co-maintain a construct of *Escherichia coli* harbouring a non-transmissible plasmid, with a tetracycline resistance gene, in competition assays without using antibiotics. Moreover, the bacterium harbouring the plasmid preserved it, without exposure to tetracycline and despite a 40% reduction in growth rate it imposes on the bacterium. In other words, my results suggest that selection for plasmids may be stronger than previously thought—explaining the plasmid paradox. This derives from sub-optimal predictions from the fitness metric above. So, here I present an iteration of the relative fitness (*W*_*ij*_) metric above to help predict this change, exposing a shift in weighting from a growth rate-dominated fitness to a population size-dominated fitness using data easily measurable in the laboratory.

## II. RESULTS

Let me assume two competing genotypes, mutants A and B, that grow consistently with population genetics theory [1, 2]. The following system of ordinary differential equations describes the change in the number of individuals of each mutant over time:

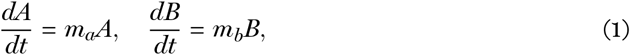

where *m* is the population growth rate increase as introduced above, and *A*(0) = *B*(0) = *x* where *x* is strictly positive. Note that *m* is derived from random birth (*b*) and death (*d*) rates, so that *m* = *b* − *d* [1], and that it is independent of the number of individuals of each genotype. Each individual is equally likely to die or reproduce at any given time, and the age distribution is near to the equilibrium so that *b* and *d* are nearly constant. The growth dynamics of both mutants is illustrated in Figure 1A. If *b* and *d* are constant, *m* is a constant and therefore the relative fitness difference between genotypes *W*_*ba*_ = *m*_*b*_/*m*_*a*_ = *k* is also constant. In other words, if 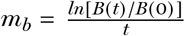 as derived from the exponential model introduced above [2, 8, 38] (see Methods), and *B*(*t*) is the number of individuals of mutant B at an arbitrary time *t*, the relative fitness *W*_*ba*_ is the same regardless of *t*. The same applies to the fitness of A with respect to B. Indeed, if *m*_*b*_ *> m*_*a*_, the number of mutant B individuals is higher than those of mutant A at all times as Figure 1A shows.

**Figure 1.**
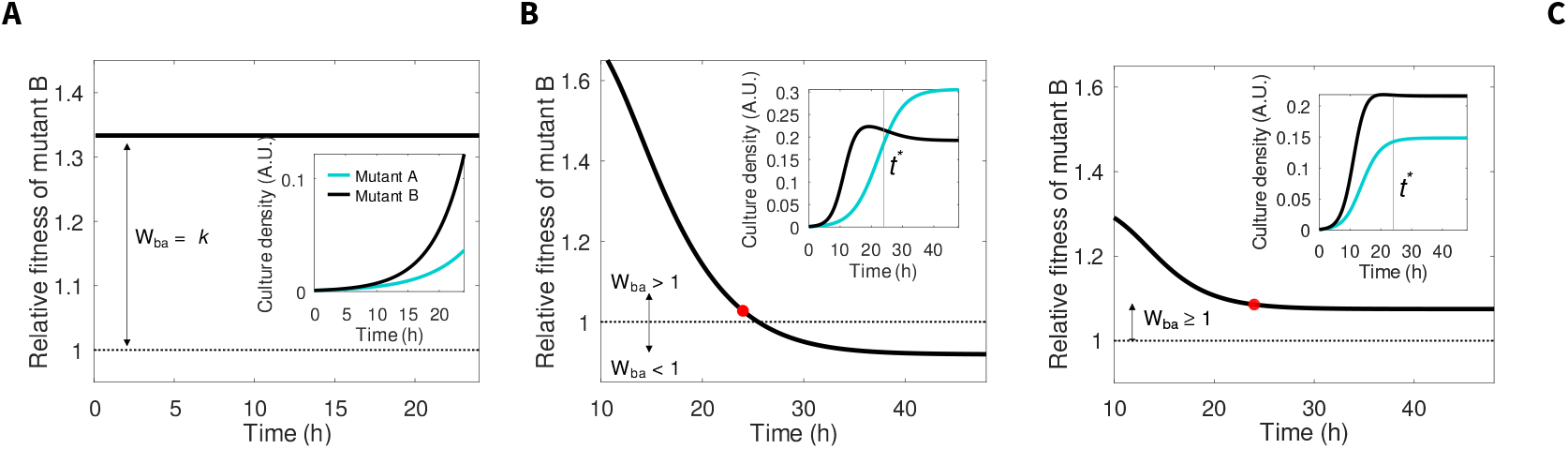
Change in relative fitness when populations have exponential and logistic growth. **A)** Change in relative fitness of mutant B (W_ba_, black) over time when competing mutants have density-independent, exponential growth—assumed by population genetics theory for populations with continuous growth and overlapping generations [1–3]. I calculated the relative fitness as [8] *W*_*ba*_ = *m*_*b*_/*m*_*a*_, where *m* corresponds to the population growth rate for each mutant. The fitness of mutant A, reference, is shown as a black, dotted line. The inset shows the change in cell density, in arbitrary units (A. U.), during the competition. **B-C)** The same information is shown for mutants with density-dependent growth, with (B) and without (C) trade-off between reproduction rate and survival. The sampling time *t*^∗^ = 24 hours, commonly used in microbiological assays, is noted by a red marker in the main plot, and a vertical, grey line in the insets. Model parameters for A) are *m*_*a*_ = 0.15 h^−1^, *m*_*b*_ = 0.2 h^−1^; for B) *m*_*a*_ = 1.0275 h^−1^, *m*_*b*_ = 1.05 h^−1^, *K*_*a*_ = 12.5 OD, *K*_*b*_ = 5 OD, *α*_*ab*_ = *α*_*ba*_ = 0.15; and for C) *m*_*a*_ = 1.0275 h^−1^, *m*_*b*_ = 1.05 h^−1^, *K*_*a*_ = 10 OD, *K*_*b*_ = 5 OD.

Now suppose that *m* depends on the number of individuals of each genotype. This is a reasonable assumption given that resources are depleted during microbial growth, and that resources are finite, limiting the abundance of each mutant over time:

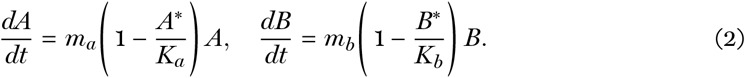

Here *K*_*a*_ and *K*_*b*_ are the maximal population size attainable (carrying capacity or population size at the equilibrium) for mutants A and B, respectively, resulting from the limited availability of resources—note different efficiencies using the *same* resource can lead to different carrying capacities [39]. Now, the population growth rate *m* is corrected by the term 1 − (*N*_*i*_/*K*_*i*_). The Lotka-Volterra model of competition [40] includes the term 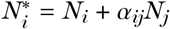, where *α*_*ij*_*N*_*j*_ is the linear reduction in growth rate—in terms of *K*—of species *i* by competing species *j*. This formalisation describes the limitation in growth imposed by the environment, due to finite resources, reducing *m* and the growth of both genotypes over time (Figure 1B and C). In this scenario, *m*_*b*_ is now corrected by changes in population size and, therefore, the above expression for *W*_*ba*_ must depend on time:

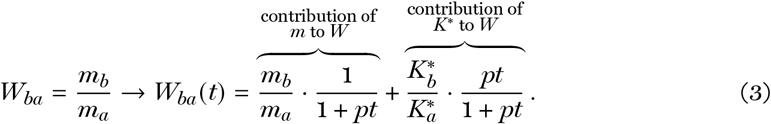

Here *W*_*ba*_ declines exponentially with time as Figures 1B and C illustrate, and eventually stabilises when the competing genotypes attain their carrying capacities *K*^∗^—new since their full potential cannot be realised in competition. Now, initially *W*_*ba*_(0) ≈ *m*_*b*_/*m*_*a*_ because 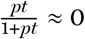 and 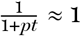, with *p* being a parameter that controls the steepness of the change. As both genotypes grow, *W*_*ba*_ declines exponentially with 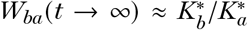 since now 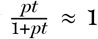 and 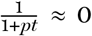. In other words, while fitness would depend on intrinsic growth rate (*m*) originally, once both genotypes are in equilibrium fitness will depend on the population size (*K*^∗^). Estimating fitness differences between two genotypes using equation (3), thus, requires prior knowledge of the carrying capacity.

This contrasts with the use of logarithms in the laboratory to estimate intrinsic growth rate as 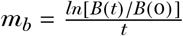 and *W*_*ba*_ = *m*_*b*_/*m*_*a*_ [8, 11, 13–16, 41–43]. Which means, there is an assumption of exponential growth as in Figure 1A. The result, however, is unreliable fitness estimates that can hinder, for example, comparative studies [44]. Figures 1B and C illustrate the importance of the sampling time for these estimates and how the use of logarithms can yield different—even conflicting—estimates. Thus, while it can be argued that estimations of *m* aim to *only* capture the exponential growth, the rest of the growth dynamics cannot be overlooked.

If, say, *m*_*b*_ *> m*_*a*_ and *W*_*ba*_ *>* 1, which mutant, A or B, would be selected in pairwise competition assays? Despite the simplicity of equations (1) and (2), the answer is not straightforward. Figure 2 illustrates the change in relative abundance of two genotypes, A, and B, growing in competition for a common resource. These genotypes are haploid, with continuous growth and overlapping generations where *m*_*a*_ and *m*_*b*_ represent their population growth rates as introduced above. The competition is propagated at regular intervals *c* where the new initial conditions are given by *N*_*a*_(*c*_+1_, 0) = *N*_*a*_(*c, t*^∗^)*d, N*_*b*_(*c*_+1_, 0) = *N*_*b*_(*c, t*^∗^)*d*, with *t*^∗^ being the time of growth allowed prior to the propagation and 0 ≤ *d* ≤ 1 the dilution factor during the propagation step analog to experimental assays [8, 12, 15]. Following population genetics theory [1, 4, 5, 8], the outcome of this competition is straightforward: if, say, mutant B reproduces faster than mutant A (*m*_*b*_ *> m*_*a*_), then *W*_*ab*_ = *m*_*a*_/*m*_*b*_ *<* 1 and mutant A goes extinct (Figure 2A). Regardless of how long the competition is allowed to progress until the next propagation step, mutant A is always lost in competition with B. Only through the emergence of a new mutant A^∗^ with 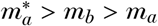 should this prediction change.

**Figure 2.**
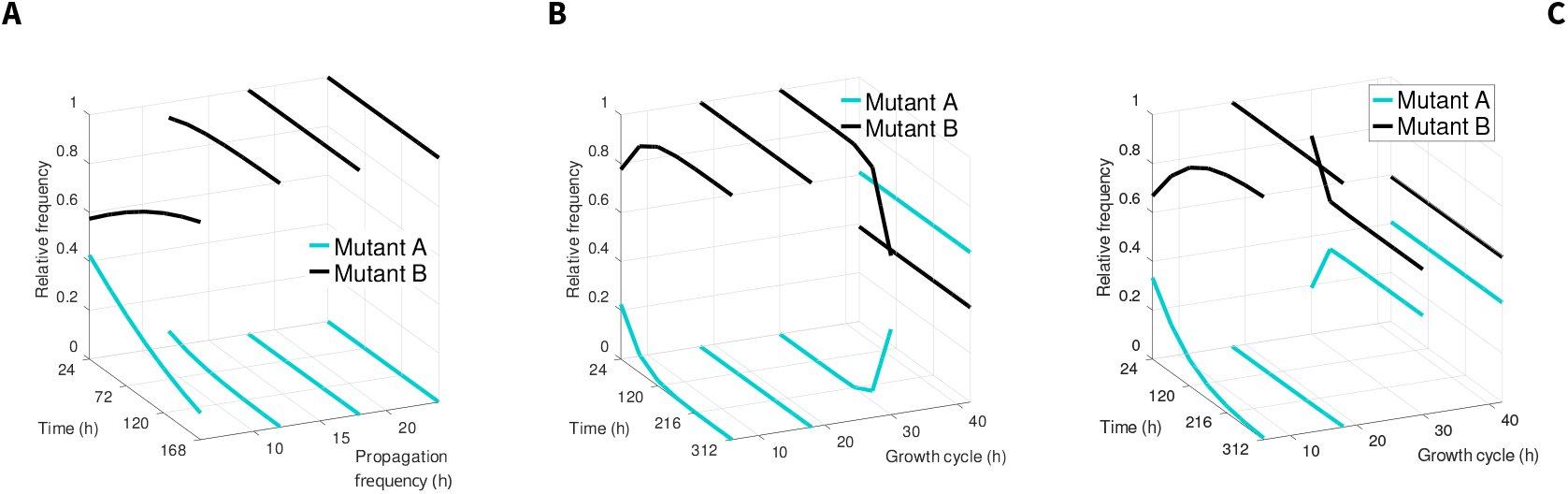
Change in relative frequency during competition depending on the length of the growth cycle. Changes in frequency over time for mutants A (cyan) and B (black) following propagations at arbitrary times t=6h, t=12h, t=18h, and t=24h. These times correspond to different stages of growth as Figure 1 shows. Each plot illustrate the case for genotypes with density-independent growth (A), or density-dependent growth where a genotype does engage in a trade-off between reproduction and survival (B) and when it does not (C). Parameter used in the model are those from Figure 1.

But when fitness changes through time the outcome of the competition becomes unclear, now depending on whether the competing genotypes reach their carrying capacities before the propagation: if the competition is propagated frequently, so that *both* genotypes are in active growth, the genotype reproducing faster will replace that reproducing slower consistently with population genetics’s notion of fitness—for this scenario. However, if one or both genotypes are allowed to reach their carrying capacities, both will be be co-maintained regardless of the in rates of population increase (Figures 2B andC). Here the relative frequency will be given by their carrying capacities *K*, so if reproduction and carrying capacity engage in a trade-off [39] for one genotype, this will be most frequent (Figure 2B). Importantly, these predictions are maintained regardless of the initial frequency of A (Figure S1), consistently with prior data [45].

To test this prediction, I competed two constructs of *Escherichia coli* MC4100, Wyl and GB(c) (see Methods), and measured their variation in fitness through time. Figure 3A shows that both constructs reach their carrying capacities within 24 hours. The construct GB(c) carries the non-transmissible plasmid pGW155B [46] harbouring *tet(36)*, a ribosome-protection type resistance gene against tetracycline [46]. This plasmid lacks a *rep* gene to control tightly the partition of plasmids after cell division [47–49] (addgene vector database accession #2853). While pGW155B is a synthetic plasmid, many natural plasmids also lack partition systems [49, 50]. Both constructs have identical chromosome with exception of this plasmid, and the fluorescence gene they carry: cyan (*cfp*, GB(c)) or yellow (*yfp*, Wyl), to allow their identification in competition assays. Using a non-transmissible plasmid prevents cross-contamination by horizontal gene transfer between the resistant construct GB(c) and Wyl. Importantly, the use of constructs derived from MC4100 avoids interactions between competitors that may affect the outcome of the competition for reasons beyond pGW155B, like the production of bacteriocins [51].

**Figure 3.**
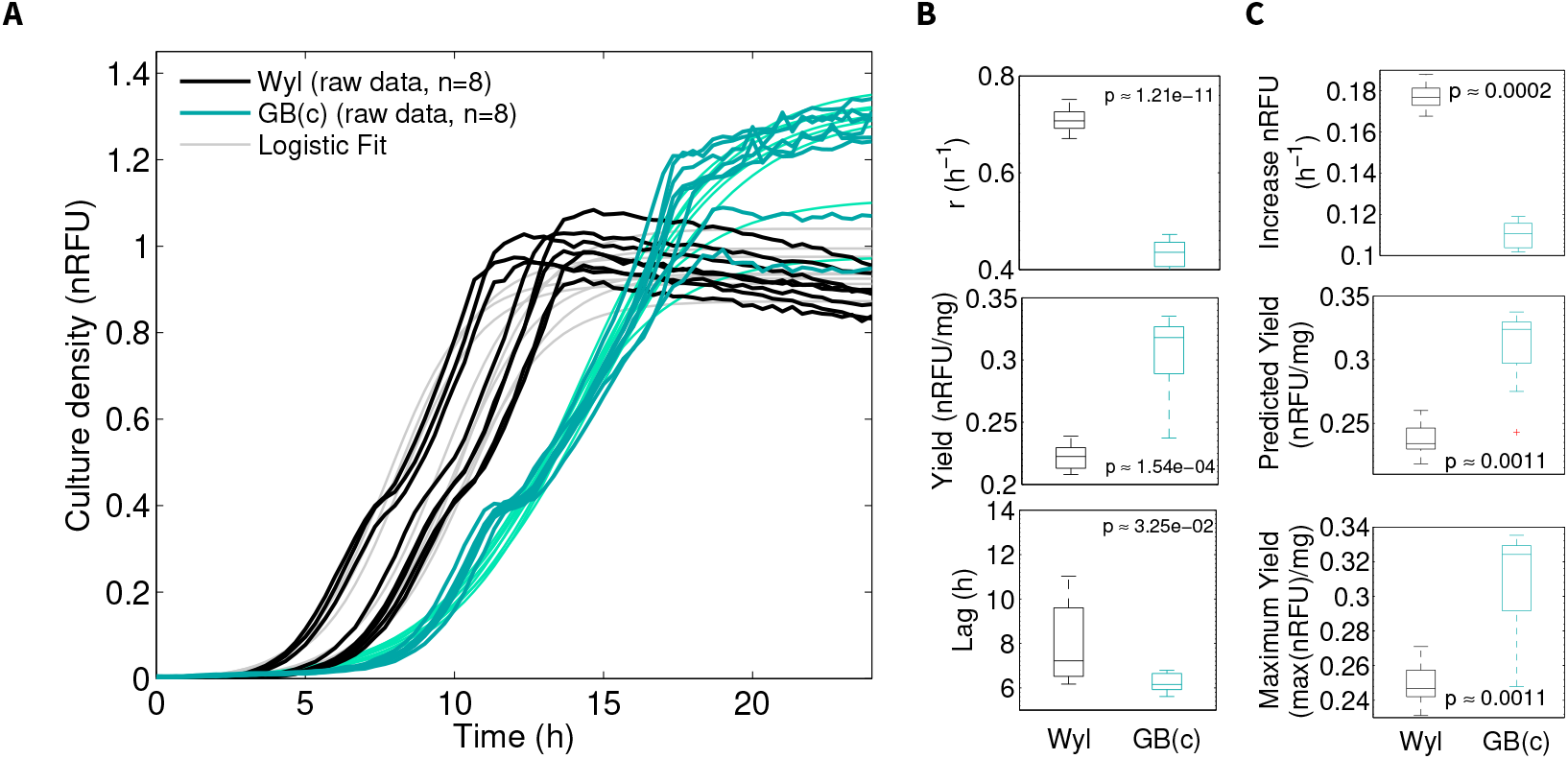
Asymmetric carriage costs of pGW155B. **A)** Overlapped growth curves of constructs Wyl (black) and GB(c) (cyan) as pure cultures, in the absence of tetracycline, over 24h. I estimated the *per capita* growth rate (*m*), population size in the equilibrium (K), biomass yield, and lag duration from a 4-parameter logistic model fitted to growth data (see Methods). Data fits for constructs Wyl and GB(c) are shown in grey and light cyan, respectively. **B)** Box plots showing the median (centre of the box), 25th and 75th percentile for each of the aforementioned parameters. The whiskers extend to the most extreme data points that are not outliers, and these are individually represented. The *p* value shown refers to a Two-sample *t*-test with unequal variance (8 replicates) that I used to test differences in the parameters between both constructs. **C)** Alternative metrics for growth rate and biomass yield: forward Euler approximation (top), data fit predicted yield (middle), and maximal yield across all time points in growth data (bottom). The *p* values correspond to Two-sample *t*-tests with unequal variance (8 replicates).

pGW155B penalised the population growth rate of construct GB(c) by approximately 40% compared to Wyl (Figure 3B) as I found out using pure cultures. It might be argued that the 4-parameter logistic model I used does not fully capture the growth data. However, if I measure the maximum change between the reads (Euler’s method, see Methods), the strong difference in growth rate is maintained. The duration of lag, and carrying capacity were also sensitive to plasmid carriage, regardless of whether I measured the growth using fluorescence or light scattering (Figure S2). Changes in cell size could confound optical density and fluorescence readings, but this phenomenon leaves a signature [52] in growth data that is absent in my dataset (Figures S2, S3). The change in carrying capacity *K* can be linked to an increase in biomass yield (*y*) as Monod’s expression [53] *y* = *K*/*S* suggests, where *S* the supply of glucose. This metric is consistent with the data, given that construct GB(c) has to express and translate plasmid-borne genes with the same supply of glucose (Figures 3B and C). This means that pGW155B triggers a metabolic trade-off between growth rate and biomass yield akin to that described when *E. coli* transitions from an inefficient to efficient metabolic pathway [39]—perhaps not surprising since the expression of plasmid-borne genes demand significant quantities of ATP [54] likely forcing cells to use respiration to yield more ATP molecules per molecule of glucose [39]. This is indeed what Figures 3B, 3C, and S4 suggest—in line with similar trade-offs described in natural plasmids [20, 55].

For the competition assay, I mixed equal proportions (cell/cell) of two pure cultures, from construct GB(c)—harbouring pGW155B—and Wyl respectively, grown overnight, in media containing no antibiotic or 0.04 µg/mL of tetracycline (see Methods). I incubated the mixed culture at 30^*o*^C until both constructs attained their population size (*K*, ∼ 24h as per data in Figure 3A), and then I propagated the competition into a new plate containing fresh media. I repeated the propagation step seven times totalling between 77 (GB(c)) and 119 (Wyl) generations given the reproduction rates in Figure 3B, where the generation time per hour is given by *m*_*i*_/24, multiplied times the number of propagation steps. Considering the mutation rate of enterobacteria [56] and the emergence of mutants after 3 growth cycles [57], I constrained the fitness measurements in both constructs over the first 24h of competition and compared the outcome of the competition with predictions in Figure 2. Indeed, the relative fitness of both constructs changed through time consistently with Figure 1B and C. In the presence of tetracycline, the relative fitness of drug-sensitive construct Wyl was below GB(c), which harbours pGW155B, at all times as Figure 4A illustrates. This meant that GB(c) increased its relative frequency in subsequent propagation steps and became the most abundant construct throughout the competition (Figure 4B) consistently with the theoretical predictions. Now, the model does not implement any mutation processes but these occur during this assay. For example, the number of copies of pGW155B harboured by GB(c) increased 5-to 6-fold during the competition (2-sample *t*-test with unequal variances, *p*=0.0088, *df*=2.5826, *t*-statistic=-7.2441; with 3 replicates) driven by exposure to tetracycline (Figure 4C) as it would be reasonable to expect given the exposure to tetracycline. Despite the occurrence of these mutations, Wyl never goes extinct consistently with Figure 2 and its proportion is determine by its abundance at the propagation time.

**Figure 4.**
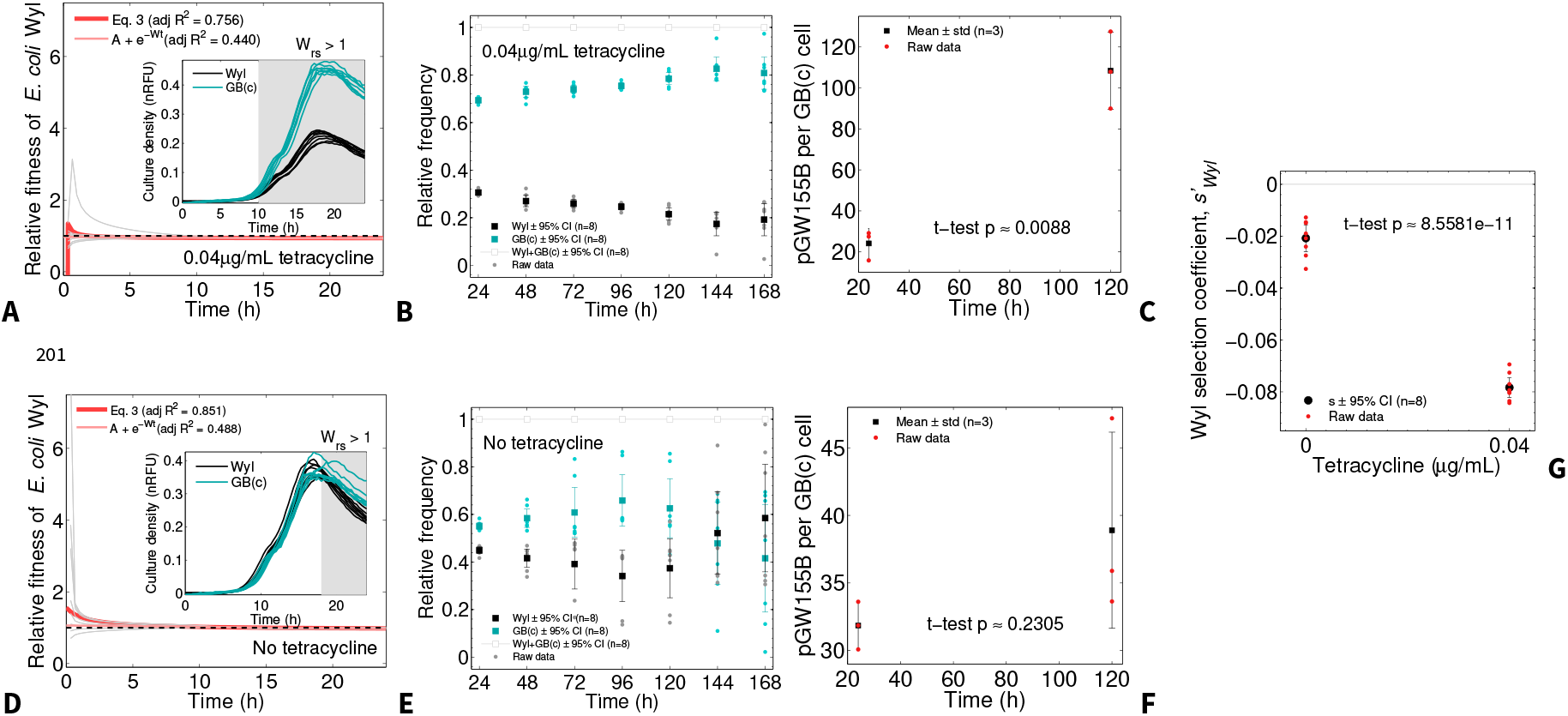
Time-dependent fitness data allows co-maintenance of both constructs without tetracycline. Relative fitness during first 24 hour pairwise-competition of construct Wyl, where both constructs grew in media supplemented with 0.04 µg/mL of tetracycline (**A**), and without antibiotic (**D**). Fitness for each replicate, derived from the logistic fit described in the Methods section as used in Figure 3A, is presented in grey with respect to the reference GB(c) (black dashed line), and the prediction from robustly fitting equation 3 to data in red with an adjusted *R*^2^ of 0.756 (A, F-statistic against zero model = 1.47 · 10^6^, *p* ≪ 10^−100^) and 0.851 (D, F-statistic against zero model = 1.12 · 10^6^, *p* ≪ 10^−100^). The inset illustrate the growth curves for both constructs, highlighting in grey the period where the relative fitness of Wyl (W_sr_) is lower than its tetracycline-resistant competitor’us. Note that the frequency of construct Wyl remains above zero at all times. **B** and **E** show the change in relative frequency of both constructs through time when both constructs grew in media supplemented with (B) and without tetracycline (E). Error bars represent the mean ± 95% confidence intervals, with raw data points shown as red dots. **C** and **F** show the change in relative copy numbers of pGW155B borne by construct GB(c) during the pairwise-competition, calculated using quantitative PCR (see Methods). Error bars represent the mean ± standard deviation, with raw data shown in red. **G** illustrates the selection coefficient for construct Wyl during the experiment (mean ± 95% confidence intervals, with raw data as red dots), a common selection metrics calculated as described in [8]. The *p*-value shown in **C, F**, and **G** corresponds to a Welch’s *t*-test—two sample *t*-test assuming that samples come from a normal distribution with unknown means, and unknown and unequal variances.

Similarly, when both constructs grew in mixed cultures, now without tetracycline exposure, the relative fitness of Wyl was higher, equal, or lower than plasmid-harbouring construct GB(c) through time, as Figure 4D shows. Consistently with the predictions both constructs where co-maintained, and with similar relative frequencies, despite the difference in growth rate (Figure 4E). Note here that while the initial deviation is small, without strong selection by antibiotic exposure each replicate evolves differently, with some replicates of Wyl being lost, while others are co-maintained. In other words, GB(c) preserved pGW155B despite the lack of tetracycline and the difference in growth rates (2-sample *t*-test with unequal variances, *p*=0.2305, *df*=2.2351, *t*-statistic=-1.6353; with 3 replicates. Figure 4F). This contrasts with the negative selection coefficient [1, 8] for construct Wyl (Figure 4G), for example, calculated as its relative fitness with respect to GB(c) measured after 24h of competition minus 1, as introduced above. This common metric suggests that Wyl is being selected against, but data show this construct is indeed being co-maintained—albeit at a lower frequency with respect to GB(c) consistently with predictions in Figure 2.

## III. DISCUSSION

The conventional notion of fitness, based on the introduced model of exponential growth, applies within a very specific set of assumptions [7]: Haploid genotypes with continuous growth and overlapping generations. Diploid genotypes or those with more complex reproductive cycles require different assumptions, and yet, due to the simplicity of the method, the same metric is used to measure fitness in organisms that range from yeast to corals [i.e. 13, 17, 58]. Chemostats [59] can closely reproduce the assumptions of exponential growth model, however it is not possible to use them more widely—i.e. in a clinical laboratory—given the complexity of the device.

Whether growth rate alone is a good proxy for fitness has been discussed at length in the literature [9, 60–62], but it is the theory that is discussed—not a method that can be implemented in the laboratory. It is noteworthy to note that fitness, taken as the aforementioned Malthusian parameter, indeed varies through evolutionary timescales due to mutations [8, 63]. My contribution here is to demonstrate that fitness *also* changes at smaller timescales, where it is assumed constant in the absence of mutations and often used to predict the fate of plasmids or indeed antibiotic-resistant mutants [11, 14–16, 20, 64, 65].

Curiously, fitness is known to be sensitive to sampling times (*t*^∗^) in the laboratory [45, 66]. But it is presented as an experimental quirk. If the sampling time *t*^∗^ is too long one competitor may drive the other to extinction, whereas if *t*^∗^ is too short the measurement may not be precise [66]. Thus, the implementation of fitness in the laboratory has been rather ‘unscientific’. As Figure 1A shows, under the exponential growth model fitness remains constant regardless of when it is measured. Now, one could argue that variations in fitness measurements are caused by frequency-dependent selection [45]. The model in [45] is based on changes in relative frequencies of two competing genotypes throughout an assay and has problems: the model has to be parameterised first, making its implementation in the laboratory somewhat problematic, and the frequencies are only measured once at the end of the assay. Fitness would still depend on one parameter only—here frequency instead of growth rate—and thus their model fits poorly the data.

The selection coefficient depends on *W*_*ij*_ since *S*_*ij*_ = *W*_*ij*_ − 1. Given the higher carrying capacity of GB(c), construct harbouring the plasmid, this metric suggests that Wyl, the competing construct, is selected against. The conclusion is therefore that Wyl will be lost to selection, but the predictions in Figures 2B and C suggests there is no such thing—outcome of competition is always co-existence, as long the competitors reach their carrying capacities. The data in Figures 4B and E show the co-maintenance of both constructs, albeit with some replicates being close to extinction—possible if a mutation emerges and the mutant no longer reaches the carrying capacity by the time at which I propagate the cultures. In other words, if relative fitness is a function of time and *W*_*ij*_ → *W*_*ij*_(*t*), the selection coefficient must also be a function of time. Whence the importance of sampling time in fitness measurements.

So, if metrics relying on growth rate are unable to capture the data, the next logic step is to introduce another trait that can be measured: population size. After all, they are two of the central parameters of the logistic growth model. Despite its simplicity, Equation 3 captures the change in weighting from a growth rate-driven fitness to a population size-driven fitness. While the metric is not perfect, since 15%–25% of the data in Figures 4A and D is not explained by the metric, it captures the essence of the transition to help make informed predictions. Despite the lack of mutations in the model, it still clearly predicts the co-maintenance of both GB(c) and Wyl—something remarkable since one would expect the tetracycline-sensitive construct Wyl to be lost.

With regards to the plasmid paradox, if the fitness measured experimentally varies within a season or cycle, it means cells harbouring plasmids—paying a cost for their maintenance—are mistakenly assumed to go extinct without positive selection. In rare cases where data is available [20, 55], it turns that when transconjugants often attain larger population sizes and the fitness measurements report a ‘carriage benefit’ despite their slower growth rates which means they will be maintained as long they reach their carrying capacity.

And this can hinder the management of antibiotic-resistance in the clinic. The fact that GB(c) maintained pGW155B even without exposure to tetracycline means antibiotic-resistant pathogens harbouring plasmids could be difficult to eradicate by simply reducing antibiotic exposure as it is commonly proposed by antibiotic stewardship guidelines.

In conclusion, growth rate-centered fitness metrics fail to capture the changes in fitness that occur at smaller timescales. These changes occur because fitness goes from being growth rate-dominated to population size-dominated, and means that a genotype can be less fit or, indeed, fitter than a reference genotype by virtue of *when* it is measured. When taking this change into account, competitions no longer have a losing side *if* both genotypes attain their carrying capacities: co-existence is the default outcome of competition, where the differences in growth rate and carrying capacities will determine the relative frequency of each competitor. When one such competitor harbours a tetracycline-resistant gene, I found the competitor is maintained in the absence of tetracycline exposure and the plasmid is preserved. This means that selection for plasmids could be higher than previously thought, explaining the plasmid paradox.

## IV. METHODS

### Media and Strains

I used the strains of *Escherichia coli* GB(c) and Wyl [67] (a gift from Remy Chait and Roy Kishony), and M9 minimal media supplemented with 0.4% glucose and 0.1% casamino acids (w/v). Note the construct Wyl harbours pCS-*λ* [68]. I made tetracycline stock solutions from powder stock (Duchefa, Ref. # 0150.0025) at 5mg/mL in 50% ethanol, filter sterilised, and stored at −20^*o*^C. Subsequent dilutions were made from this stock in supplemented M9 minimal media and kept at 4^*o*^C.

### Batch transfer protocol

I inoculated a 96-well microtitre plate containing 150 µg/mL of supplemented M9 media with a mixture of two overnight cultures, one of *E. coli* GB(c) and another of *E. coli* Wyl (1µL containing approx. 2 × 10^6^ cells, Figure S5). To avoid unforeseen variation in the initial number of copies of pGW155B between assays, the overnight culture for GB(c) was supplemented with 100ng/mL of tetracycline as described elsewhere [12]. Then, I centrifuged the cultures, and resuspended the pelleted cells in M9 minimal media prior adding to the microtitre plate to avoid the introduction of residual antibiotic and metabolic byproducts. I incubated the plate at 30^*o*^C in a commercial spectrophotometer and measured the optical density of each well at 600nm (OD_600_), yellow florescence for the Wyl strain (YFP excitation at 505nm, emission at 540nm), and cyan fluorescence for the GB(c) strain (CFP at 430nm/480nm) every 20min for 24h. After each day I transferred 1.5µL of each well, using a 96-well pin replicator, into a new microtitre plate containing fresh growth medium and tetracycline.

The use of by-hand laboratory techniques, such as colony counting, were impractical given the continuous measurements required for the experiments in this study and are also error-prone [69]. Thus, given the existing correlation between cell number, optical density, and relative fluorescence measurements (Supplementary Figures S5 and S6); I used a spectrophotometer to automatically produce proxies of population density. While these devices have inherent limitations, Supplementary Figure S5 shows that OD_600_ and fluorescence readings are reasonable indicators of bacterial population density: Readings correlate positively, and indeed linearly, with live cell counts measured in colony-forming units (CFU) per ml.

### Growth parameter estimation

Yellow and cyan fluorescence protein genes were constitutively expressed given the approximately constant ratio between fluorescence and optical density (Figure S6) in the conditions tested in the article. This allowed me to use fluorescence data as a proxy for cell density in mixed culture conditions. I normalised fluorescence readings using a conversion factor, *n*_*f*_, calculated by growing each construct in a range of glucose concentrations, and regressing the linear model RFU = *n*_f_ · OD + *c*, where RFU is relative fluorescence units data, OD optical density data, *n*_*f*_ the conversion factor between fluorescence and optical density, and *c* the crossing point with the *y*–axis when OD = 0. I imported the resulting time series data set (Figures S3) into MATLAB R2014b to subtract background and calculate fitness as described in the main text as follows.

Once corrected, I used the expression 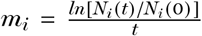 to estimate the *per capita* growth rate *m*_*i*_ as per in [8, 11, 14–16] although I used different values for *t* to test the theoretical predictions. This expression derives from the exponential model in equation 1 as it follows. We can isolate the term with *m* for, say, mutant A, as 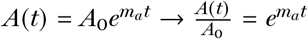. To remove the number *e* we then apply logarithms to this expression leading to 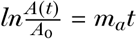. Re-arranging the terms we are left with 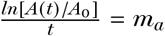, which is the expression used by experimentalists.

To measure the carriage costs of pGW155B, I fitted to data the logistic model

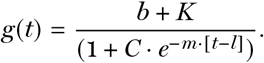

where the term *g*(*t*) denotes the change in culture growth as a function of time (in OD, YFP, or CFP units), *b* the inoculum size used to subtract the background, *C* is a parameter, *m* the *per capita* growth rate, *K* the maximal population size attained and *l* the duration of the lag phase. Alternatively, I measured the rate of increase using the forward Euler approximation 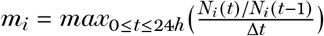, where Δ*t* = 20min = 1/3 is the time between reads. To estimate the biomass yield I divided OD data in stationary phase by the glucose supplied [70]. Alternatively, I used the highest density across all time points and the density reported by the data fit, both divided by glucose supply, as alternative metrics for biomass yield.

Finally, I divided the length of the growth cycle (24h) by the *per capita* growth rate, multiplied by the number of propagations (seven) to estimate the total number of duplication events—where one duplication means one generation—for each construct.

### DNA material extraction

For each concentration, I sampled three representative 150 µg/mL replicates that I divided in two groups: for chromosome and plasmid DNA extraction. I used ‘GeneJet DNA’ (ThermoScientific, Ref. #K0729) and ‘GeneJet Plasmid’ (ThermoScientific, Ref. #K0502) extraction kits to extract chromosomal and plasmid DNA (pDNA) from the samples, respectively, and used Qubit to quantify DNA and pDNA yields. Both extracts were diluted accordingly in extraction buffer, as per in manufacturer instructions, to normalise DNA across samples.

### Quantitative PCR and plasmid copy number estimation

I used primer3 to design two pairs of primers with melting temperature (T_*m*_) of 60^*o*^C and non-overlapping probes with T_*m*_ of 70^*o*^C. The amplicon ranges between 100 to 141bp depending on the locus (Table 1). Two reaction mixes were prepared using the kit ‘Luminaris Color Probe Low ROX’ (ThermoScientific, Ref. #K0342), adding 0.3µM of each primer and 0.2µM of the probe as per manufacturer specifications. Following a calibration curve for each reaction (Figure S7) I added 0.01ng of chromosomal or plasmid DNA material to each of the reaction mixes.

**Table 1.**
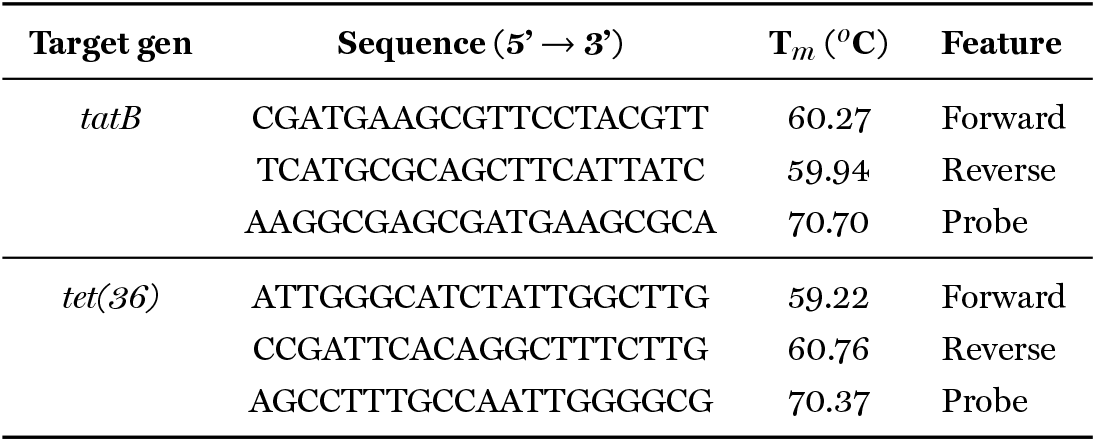
Primers and probes designed using Primer3. Amplicon ranging from 100 to 141bp. T_*m*_ indicates the estimated melting temperature.

To estimate the relative copies of pGW155B per GB(c) cell, I calculated the corresponding proportion of chromosomal DNA corresponding to the GB(c)-type from data in Figure 4 and used the expression [15]

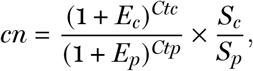

where *cn* is the number of plasmid copies per chromosome, *S*_*c*_ and *S*_*p*_ are the size of the chromosome and pGW155B amplicon in bp, *E*_*c*_ and *E*_*p*_ the efficiency of the qPCR taken from data in Figure S7, and *Ctc* and *Ctp* are the cycles at which I first detected product amplification (*C*_*t*_).

## Availability of data and materials

Data used to generate Figures 3 and 4 is available as a supplementary table. An Octave/MATLAB implementation of equations 1 and 2 is available at https://gitlab.com/rc-reding/papers/-/tree/master/FisherTheorem.

## Acknowledgements

The author thanks Robert Beardmore for laboratory support during the execution of this project.

---

**Figure S1.**
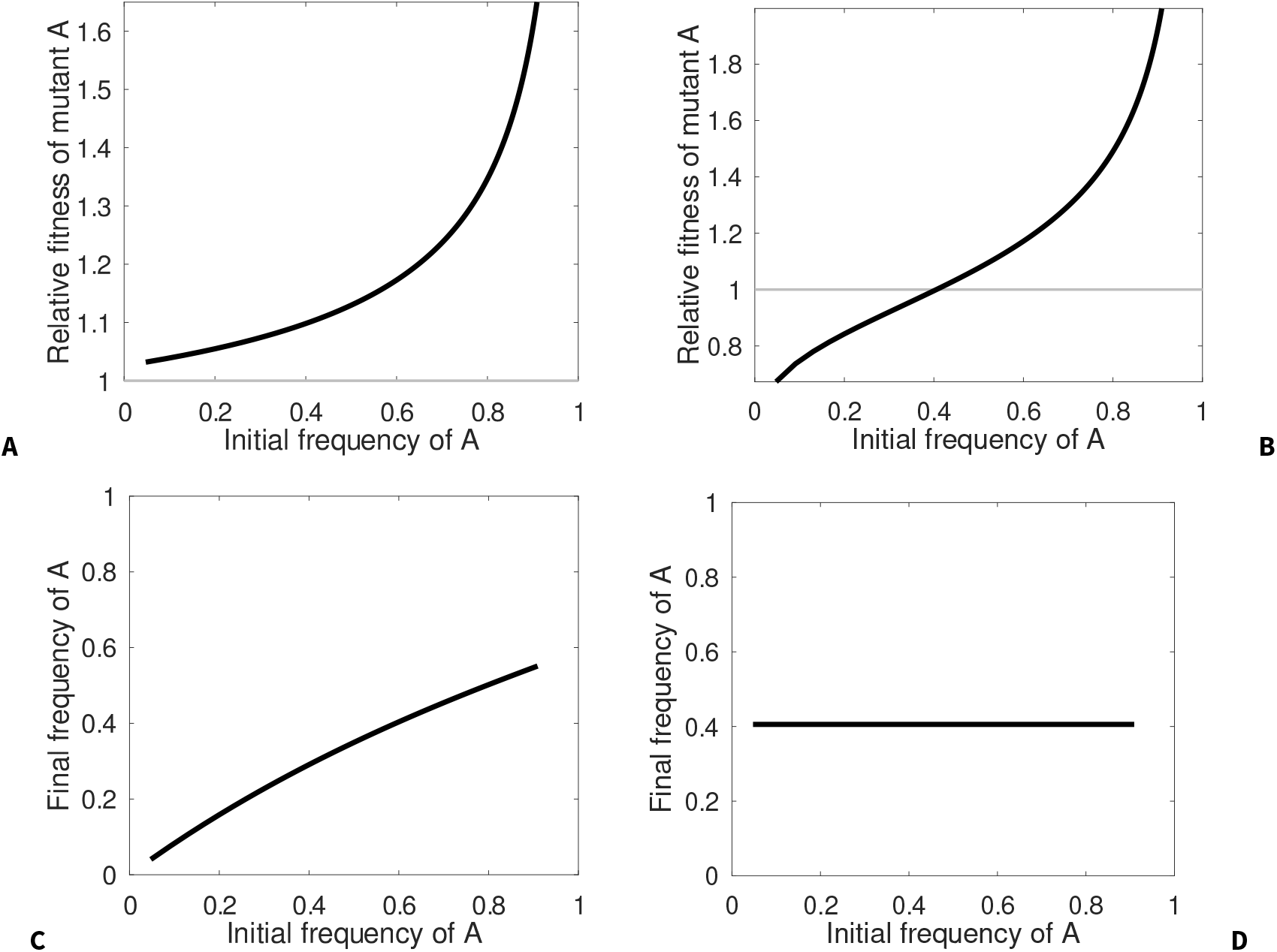
Relative fitness of mutant A at different initial frequencies. **A–B)** Relative fitness of mutant A measured after seven propagation steps, analog to those described in the main manuscript, when growth rate and population size engage in a trade-off (A) and when they do not (B). The frequency on the x-axis is that of mutant A at t=0. The fitness of each competitor (*m*_*i*_) was calculated as explained in the main manuscript as 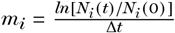 where *N*_*i*_ (*t*) is the density of each mutant at time *t* = 7, *N*_*i*_ (0) that at time t=0, and Δ*t* = 7. This was used to calculate the relative fitness of A as *W*_*ab*_ = *m*_*a*_/*m*_*b*_ as described in the text. The code is available in github in the link provided in the text, and the parameters used are those from Figure 1B and C. **C–D)** Relative frequency of mutant A at the end of the seventh propagation when growth rate and population size engage in a trade-off (C) and when they do not (D), shown in the y-axis, as a function of its frequency at t=0.

**Figure S2.**
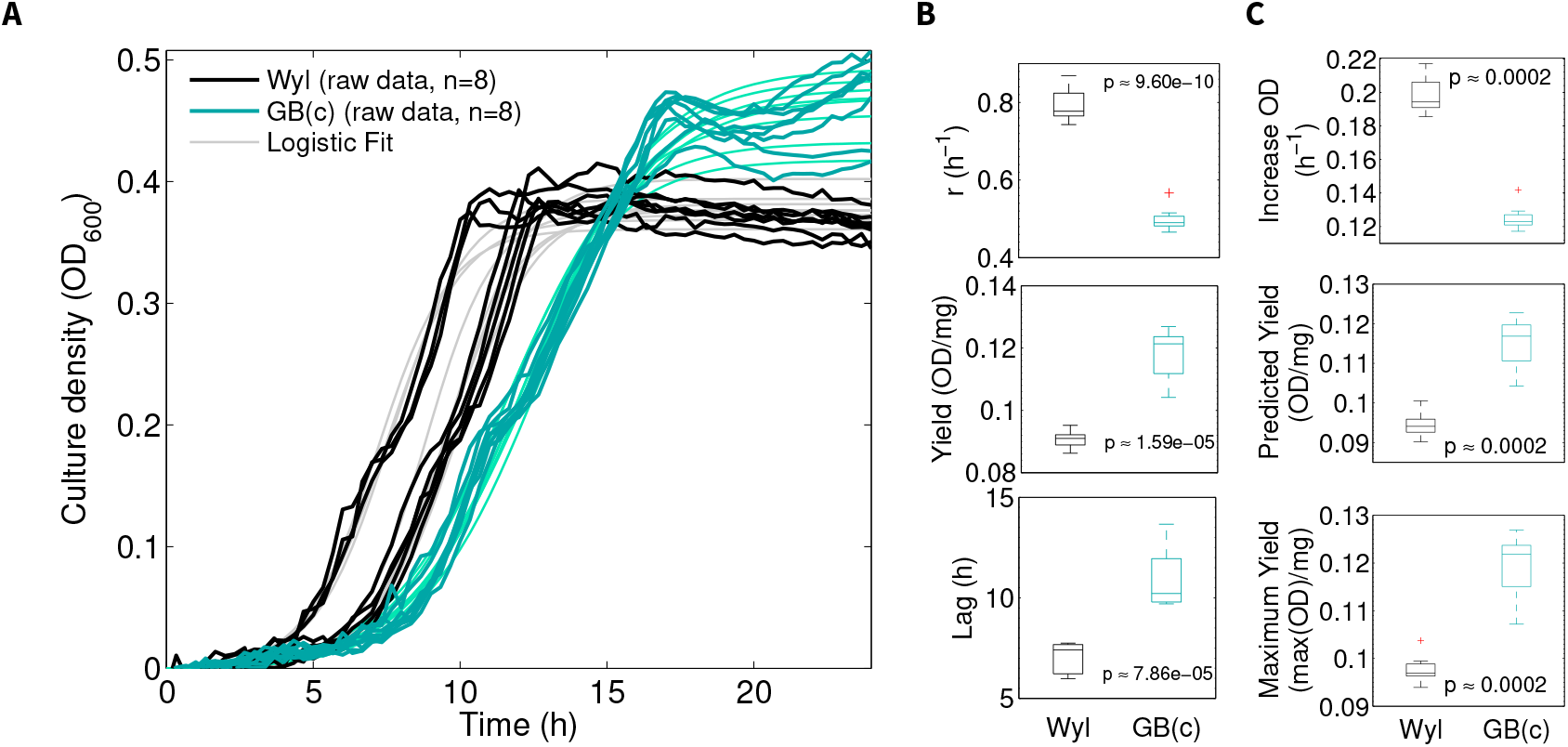
Asymmetric carriage costs of pGW155B (optical density data). **A)** Growth curves of constructs Wyl (black) and GB(c) (cyan) in pure culture, in the absence of tetracycline, over 24h. I estimated the maximum growth rate (r), population size in the equilibrium (K), biomass yield, and lag duration from a 4-parameter logistic model fitted to growth data (see Methods). Data fits for constructs Wyl and GB(c) are shown in grey and light cyan, respectively. **B)** Box plots showing the median (centre of the box), 25th and 75th percentile for each of the aforementioned parameters. The whiskers extend to the most extreme data points that are not outliers, and these are individually represented. The *p* value shown refers to Two-sample *t*-tests with unequal variance (8 replicates) that I used to test differences in the parameters between both constructs. **C)** Alternative metrics for growth rate and biomass yield: forward Euler approximation (top), data fit predicted yield (middle), and maximal yield across all time points (bottom). The *p* values correspond to Two-sample *t*-tests with unequal variance (8 replicates).

**Figure S3.**
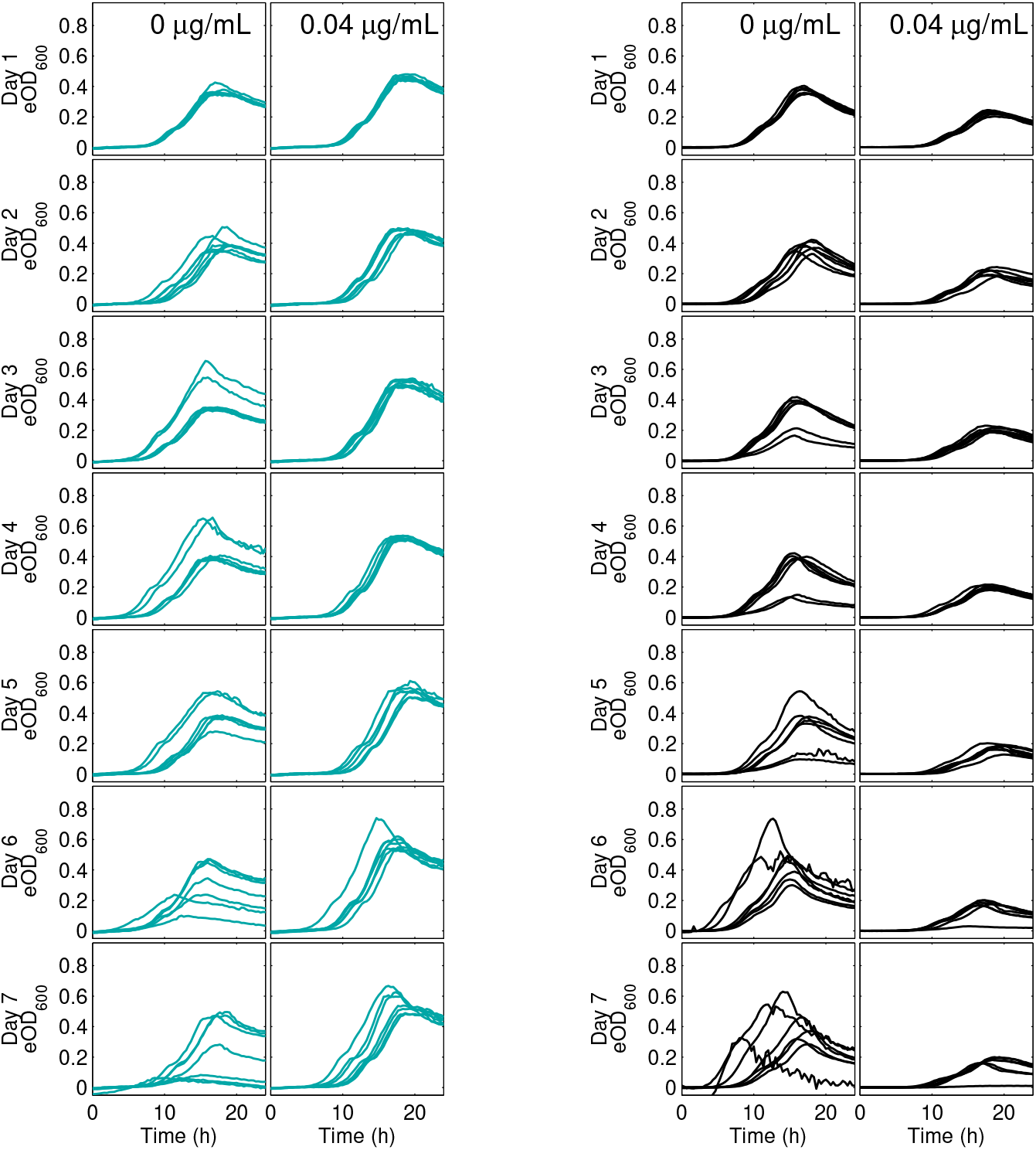
Raw data of each construct during the competition. Raw optical density data for GB(c) (cyan) and Wyl (black), measured every 20 min and converted from fluorescence readings. Each individual plot represents the change in abundance of each construct over 24 hours. Data corresponding to each condition, with and without tetracycline, are aggregated as columns.

**Figure S4.**
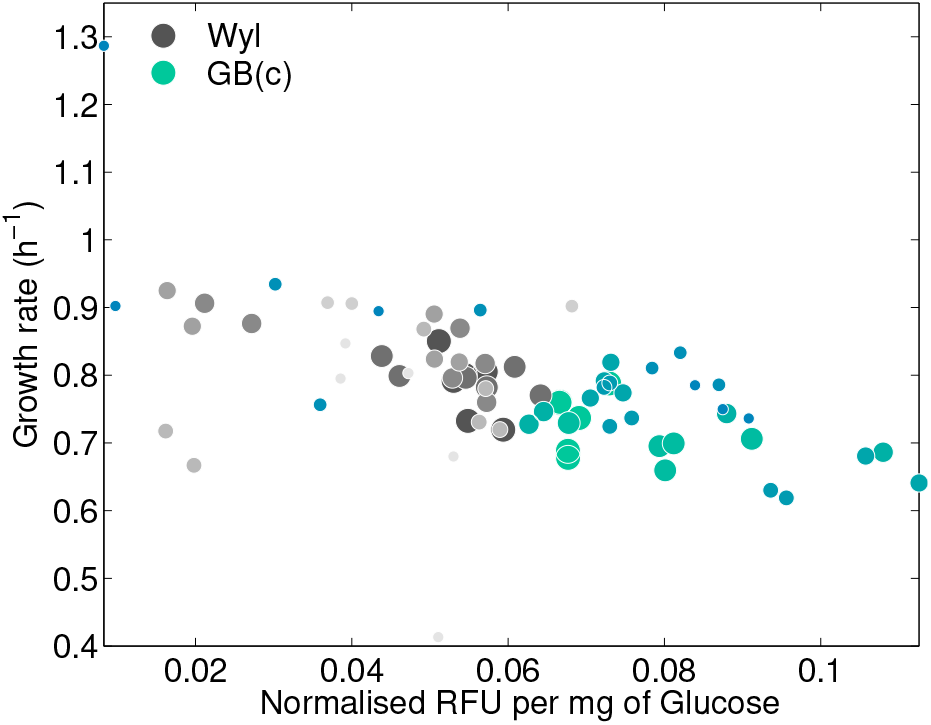
Rate-Yield Trade-Off imposed by pGW155B on construct GB(c) also during the competition. Pooled relationship between *per capita* growth rate, on the y-axis, and biomass yield during the competition assay. The growth rate parameter, as well as the carrying capacity (*K*), were estimated using the logistic model described in the methods. Biomass yield was then calculated by dividing the carrying capacity by the glucose supplied so that *c* = *K*/*Glc* [39, 70]. The data corresponding to the first day of the assays is represented with the largest dot size, with the following days having smaller dot sizes—smallest represents the data from last day of the assay.

**Figure S5.**
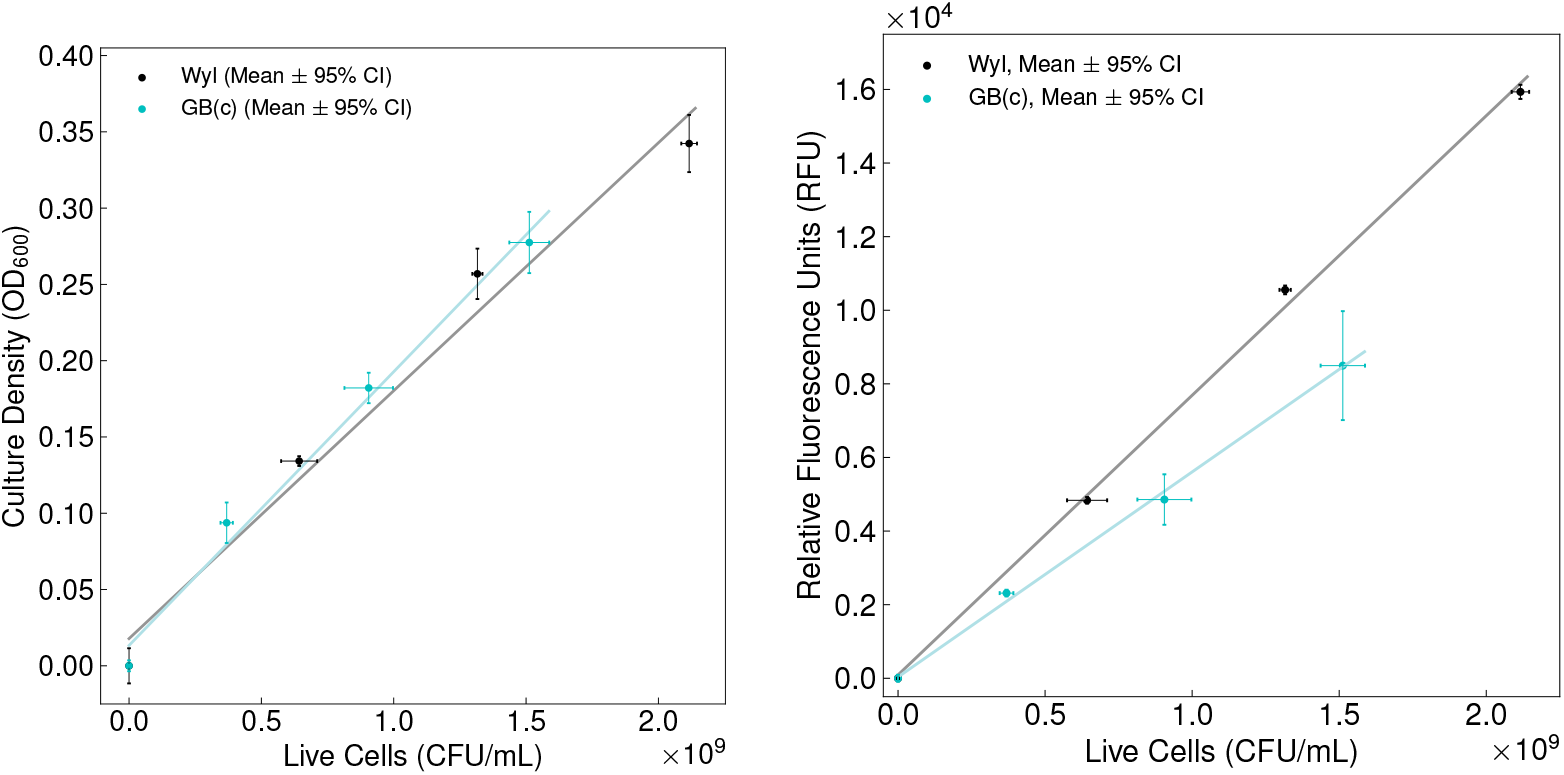
Calibration curve to translate optical density and fluorescence data to number of *Escherichia coli* cells. I fitted the linear model *a* = *bx* + *c* to optical density and colony counting data (dots) to calculate the number of optical density units (OD_600_, left) and relative fluorescence units (RFU, right) per cell. *a* denotes the optical density readings measured at 600nm, *c* the crossing point with the *y*−axis when *x* = 0, and *b* the conversion factor between optical density and number of cells (*x*). I interpolated optical density and fluorescence readings to calculate the number of cells within a culture as *x* = (*a* − *c*)/*b*. For the strain Wyl, *b* = 1.62 × 10^−10^ *OD* · *mL* · *CFU*^−1^ and *c* = 1.78 × 10^−2^ *OD*, whereas for GB(c) *b* = 1.79 × 10^−10^ *OD* · *mL* · *CFU*^−1^ and *c* = 1.33 ×10^−2^ *OD*. For the case of relative fluorescence units, *b* = 7.597 ×10^−6^ *RFU* · *mL*· *CFU*^−1^ and *c* = 90.891 *RFU* for strain Wyl; whereas *b* = 5.573 × 10^−6^ *RFU* · *mL* · *CFU*^−1^ and *c* = 33.706 *RFU* for GB(c).

**Figure S6.**
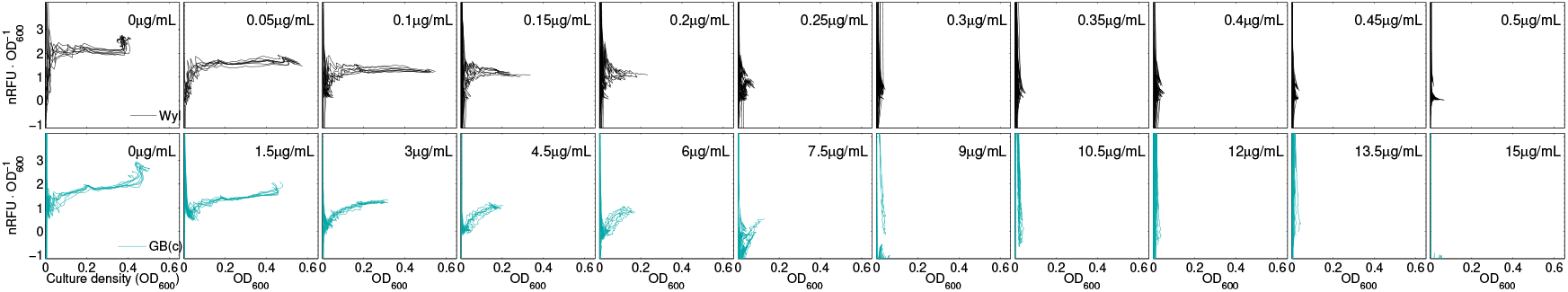
Changes in relative fluorescence over time in GB(c) and Wyl strains in pure culture conditions. Raw change in florescence, per optical density units, measured every 20min for 24h for the Wyl-(black) and GB(c)-type. Each column represents the data set for each tetracycline concentration used.

**Figure S7.**
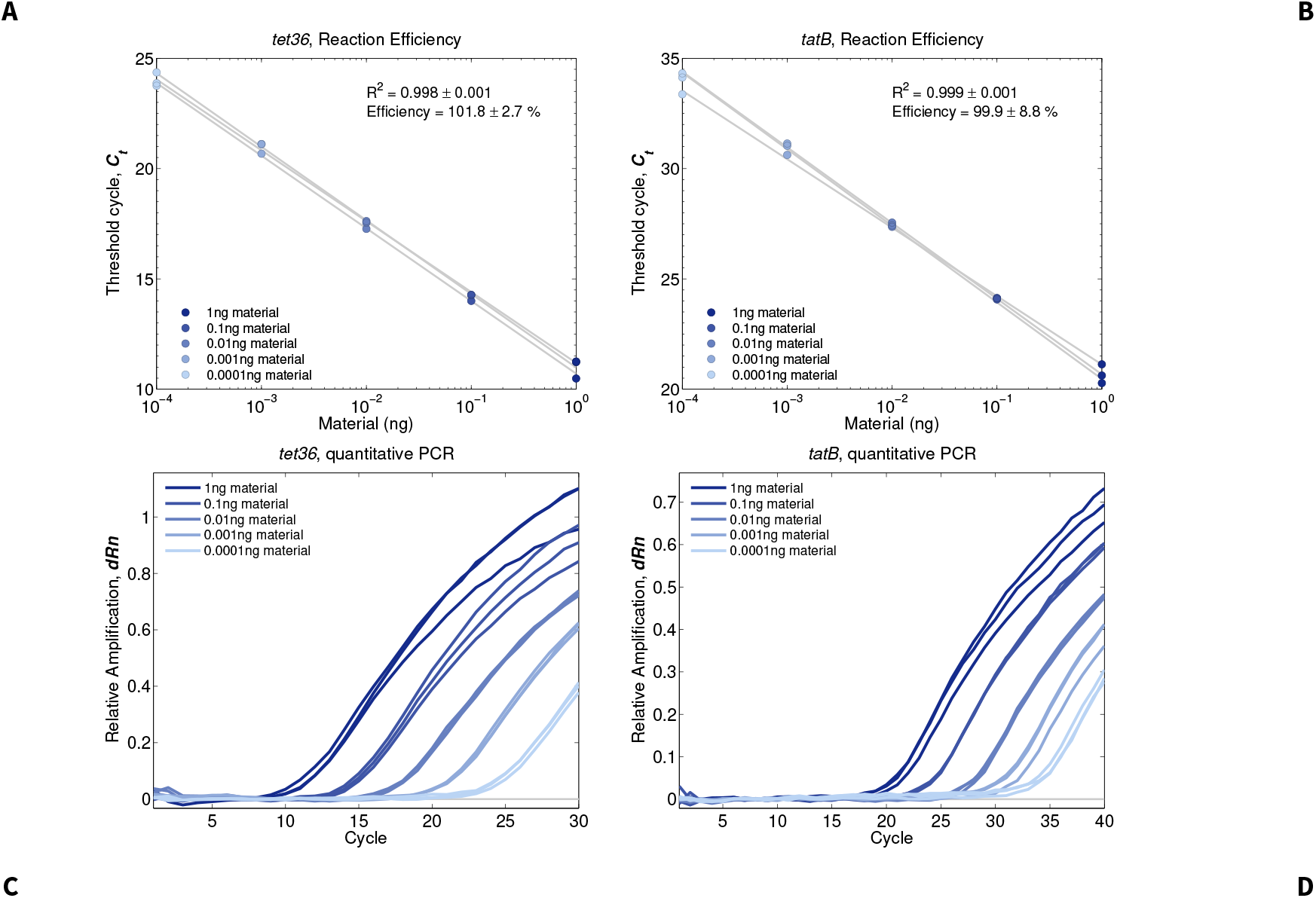
Quantitative PCR calibration curves for *tet(36)* and *tatB*. Reaction efficiency for the set of primers and probes, listed in Table 1 the main Methods section, for *tet(36)* (A) and *tatB*. The efficiency was calculated as *E*_*f*_ = 10^−1/*Slope*^ − 1, and the slope term calculated by fitting a linear model to qPCR threshold cycle (*C*_*t*_) data. The mean ± standard deviation for the adjusted coefficient of determination *R*^2^ and efficiency are shown in the figures. The amplification curves for each reaction, using 3 replicates, are shown in C) and D), respectively.

